# Community structure determines the predictability of population collapse

**DOI:** 10.1101/2022.03.07.483267

**Authors:** Gaurav Baruah, Arpat Ozgul, Christopher F. Clements

## Abstract

1. Early warning signals (EWS) are phenomenological tools that have been proposed as predictors of the collapse of biological systems. Whilst a growing body of work has shown the utility of EWS based on either statistic derived from abundance data or shifts in phenotypic traits such as body size, so far this work has largely focused on single species populations.
2. However, in order to predict reliably the future state of ecological systems which inherently could consist of multiple species, understanding how reliable such signals are in a community context is critical.
3. Here, reconciling quantitative genetics and Lotka-Volterra equations which allow us to track both abundance and mean traits, we simulate the collapse of populations embedded in mutualistic and multi-trophic predator-prey communities. Using these simulations and warning signals derived from both population- and community-level data, we show that the utility of abundance-based EWS as well as metrics derived from stability-landscape theory (e.g., width and depth of the basin of attraction) are fundamentally linked, and thus the depth and width of the stability landscape could be used to identify which species should exhibit the strongest EWS of collapse.
4. The probability a species displays both trait and abundance based EWS is dependent on its position in a community, with some species able to act as indicators species. In addition, our results also demonstrate that in general trait-based EWS appear less reliable in comparison to abundance-based EWS in forecasting species collapses in our simulated communities. Furthermore, community-level abundance-based EWS were fairly reliable in comparison to their species-level counterparts in forecasting species level collapses.
5. Our study suggests a holistic framework that combines abundance-based EWS and metrics derived from stability-landscape theory that may help in forecasting species loss in a community context.

## Introduction

Regime shifts in ecological systems are characterized by rapid transition from one stable state to another state in response to an environmental perturbation (Scheffer et al. 2009, Arani et al. 2021). There has been growing interest in predicting such critical transitions using statistical indicators, commonly known as ‘early warning signals’ (EWS) detected in time series data (Scheffer et al. 2009, Drake and Griffen 2010, Dakos et al. 2014, Baruah et al. 2020). These are phenomenological signals that are derived from the theory of alternative stable states and can be easily estimated from abundance or biomass time series data (Dakos et al. 2012). As an ecological system gets increasingly sensitive to external perturbation – and thus is at increasing risk of shifting to an alternative stable state (such as population extinction) – metrics such as autocorrelation and variance of a time series are predicted to steadily increase (Scheffer et al. 2009, Drake and Griffen 2010, Lade and Gross 2012, Clements and Ozgul 2018). Whilst much work has sought to detect such signals, the efficacy of such abundance-based signals has been called into question (Boerlijst et al. 2013, Clements et al. 2015, Dutta et al. 2018, Arkilanian et al. 2020, Baruah et al. 2021). Consequently, in order to improve forecasts of population collapse, warning signals that incorporate information from fitness-related traits (known as trait-based EWS), such as body size, have been proposed (Clements and Ozgul 2016, Clements et al. 2017, Baruah et al. 2019, 2021). Although such EWSs have been shown to forecast rapid shifts in a variety of ecological systems in response to continuous (Drake and Griffen 2010, Suweis and D’Odorico 2014) or seasonal environmental perturbation (Burant et al. 2019, 2021), relatively few studies have evaluated the performance of EWS in predicting population level collapses in a multispecies context (Boerlijst et al. 2013, Dakos and Bascompte 2014, Dakos 2017). Whilst it is known that EWS could be useful in forecasting community level transitions (Carpenter et al. 2011, Spanbauer et al. 2016), the impact of community structure and species interaction on the utility of EWS is relatively still unknown.

There is ample evidence to suggest that different species interactions and interaction type could be a key determinant of the stability of a system (Mougi and Kondoh 2012, Hastings et al. 2018), and understanding how the structure of ecological communities might alter the detectability of their collapse has obvious conservation and management implications. Species are embedded in a network of different interactions spanning mutualism, competition and predator-prey. In addition to these interactions, network architecture defined by connectance, nestedness, or link density further impacts how species interacts and maintain diversity (Dunne and Williams 2009, Landi et al. 2018). For instance, nestedness has been suggested to lead to abrupt community collapses in mutualistic networks (Lever et al. 2020, Baruah 2022). Before one is able to understand and evaluate the impact of network architecture on predictability of species or community collapses, one has to understand how type of interaction within a community could impact detectability of biodiversity collapses. Changes in the external environment could not only trigger changes in abundance but also could directly or indirectly influence changes in mean traits over time (Cortez 2016). Such changes in mean traits could be through rapid evolution thereby either delaying a transition to collapse (Dakos et al. 2019) or could also lead to and earlier collapse, such as those observed in the case of “evolutionary suicide” (Rankin and López-Sepulcre 2005). Whenever transitions to alternative stable states have been investigated in ecological systems, the evolutionary aspect of trait change has usually been overlooked. For instance, in a mutualistic community such as a plant-pollinator system, trait matching with other species can have positive consequences for the community stability as well as for species persistence in response to a perturbation (Guimarães et al. 2007). As a result of trait matching, evolutionary shifts in traits of both the plant and the pollinator could occur in a similar direction when, for instance, only one of the two species engaging in mutualistic interaction is forced to collapse. In a predator-prey community, however, shifts in resource abundance could lead to shifts in predator traits that might concurrently impact the whole community (Haney & Sipeliski 2018). These environmentally driven but interspecifically mediated trait shifts might feedback and alter abundances of other species within the network, with the type of interaction (mutualistic, or predator-prey) determining the strength and direction of shifts in both traits and abundances. Critically, in the context of EWS, this would mean that when one perturbs a single species in an ecological community the perturbation could propagate and amplify to affect whole community stability.

Thus, the strength and type of interspecific interactions might drive the direction and magnitude of trait and abundance shifts in the face of environmental change (Haney and Sipelski 2018), leading to potentially non-intuitive trends in both the traits and abundances of species interacting in a community where components (species) collapse due to external perturbation. This raises a key question in predicting the fate of biological communities: what should we monitor to allow us to best predict the future state of this system – individual species (Dai et al. 2012, Kéfi et al. 2013) or the entire community (Dakos and Bascompte 2014)? Previous EWS research has typically focused on the species of interest (Clements *et al* 2019), however alternative approaches include combining all species level data together and analyzing this for EWS (Carpenter et al. 2011), or potentially seeking ‘indicator species’ (e.g., keystone species) (Doncaster et al. 2016) which may due to their location within a food-web provide reliable warning signals of approaching collapse.

Here we explore these key issues by combining quantitative genetics and generalized Lotka-Volterra equations. Using two common ecological communities (mutualism and predator-prey) we investigate how the interaction type of a community can alter the stability and predictability of collapse. Using data at both the species and community level, we first evaluate how stability-resilience metrics derived from stability-landscape curves (see section 2.3) links with the suite of classical EWS methods, and shifts in mean trait values, as well as another warning signal designed to accommodate the high-dimensional nature of biological communities based on a shift in the dominant eigenvalue of the system (see Chen et al 2019). Forcing each species in a network to collapse independently of the others, we show that some species are more likely to display both trait and abundance-based EWS, and thus act as indicator species for the collapse of a community. In addition, our results show that the identity of such indicator species depends on the strength of interaction and overall fitness benefits gained in a community which consequently depends on the community type they are embedded in.

## 2. Methods

### 2.1 General modelling framework

We used quantitative genetics and generalized Lotka-Volterra equations to model trait-mediated ecological interactions for two community motifs, each including six-species and comprising of predator-prey interactions and mutualistic interactions (Fig. 1). Eco-evolutionary dynamics and ecological interactions were assumed to depend solely on a species’ trait in the trait axis and the distance of the species trait value from other species in the community, as given by:

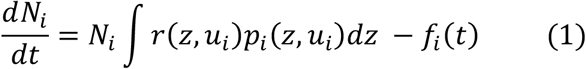

where *r*(*z*, *u_i_*) is the per capita growth rate of species *i* which is dependent on the type of interaction that is being modelled and *p_i_*(*z*, *u_i_*) is the distribution of trait *z* with mean *u_i_* and variance 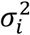. *f_i_*(*t*) is the fold harvesting function given as 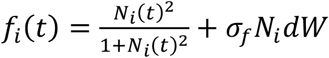 (Dakos et al. 2012), where *dW* is a white noise process with mean 0 and variance, 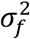 of 0.1. Eco-evolutionary dynamics of the mean trait *u_i_* in response to selection due to inter and intraspecific interactions with other species can be defined as:

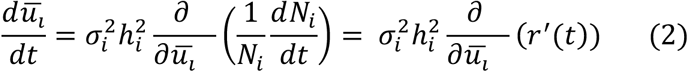

where,

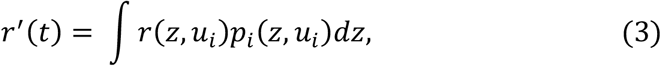

and 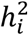 is the broad-sense heritability of the mean trait. *r*’(*t*) will differ with the type of community. All community motifs included competition among the basal species. We specifically ensure that competitive interactions are fixed (trait-independent), with intraspecific competition (*α_ii_*) being substantially higher than interspecific interactions (*α_ij_*). This specific parameterization ensures first, the stability of the community (Barabás et al. 2016, 2017), and second, that eco-evolutionary dynamics are primarily dependent on trait-mediated interactions in the two community motifs (Fig. 1). It should be noted that in our modelling framework we chose two different community motifs that are dominated by either mutualistic trait-based interaction or predator-prey trait-based interaction. In reality, ecological communities are complex and typically consist of a variety of interactions which may promote stability (Mougi and Kondoh 2012). However, rather than using a continuum of potential network types, we assess how two types of interspecific interactions (mutualism and prey-predation) affect the presence of warning signals by simulating communities whose dynamics are predominantly shaped by these interactions.

**Figure 1.**
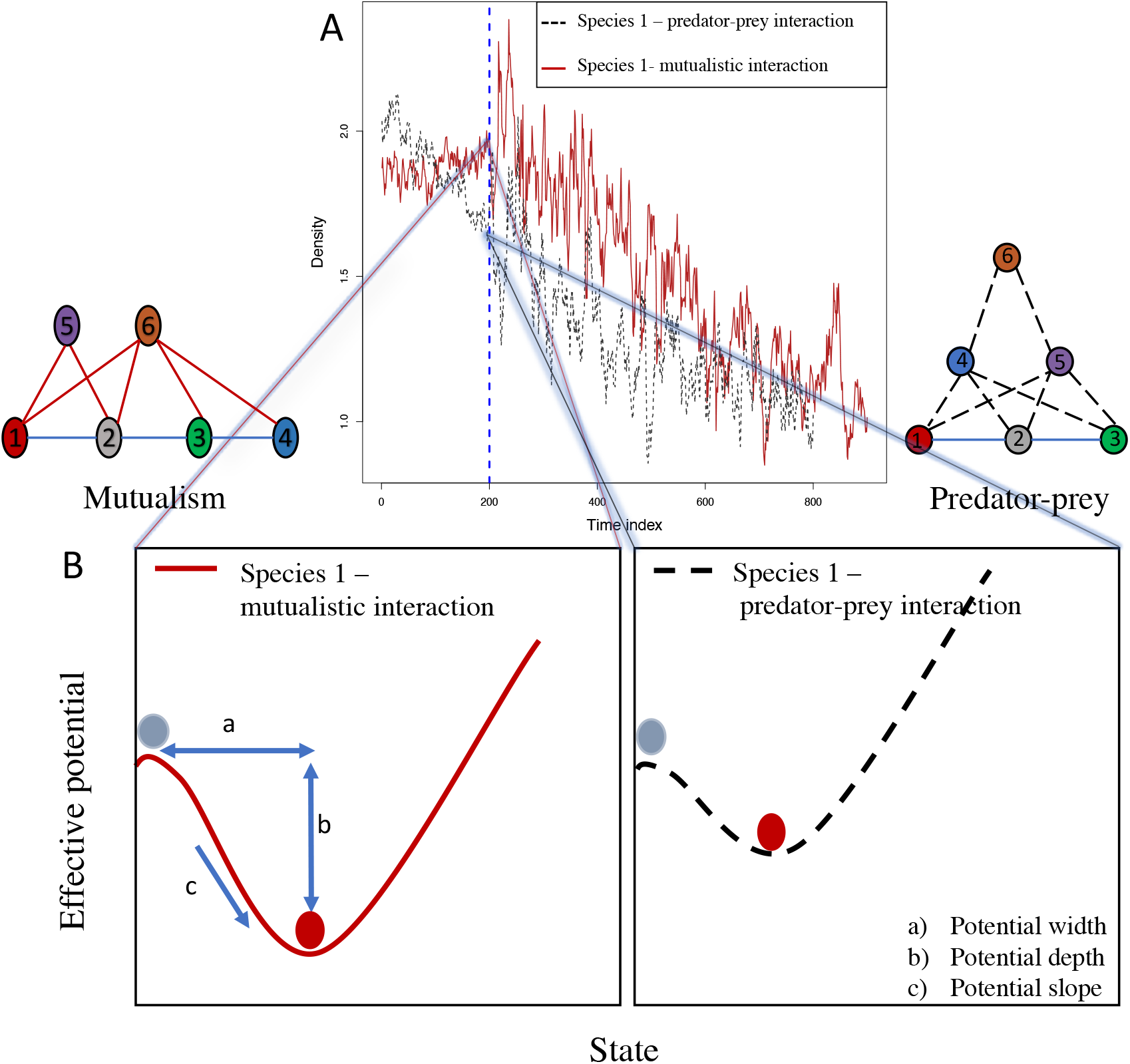
We investigate the presence of early warning signals prior to the collapse of populations embedded in two distinct community motifs: predator-prey food-web and mutualism community. A) Before the state of perturbation of a target species we quantified the effective potential curves, and three other metrics that measure resilience and stability directly from these curves. To be noted, that although resilience and stability are related, they however mean different aspects of a dynamical system. Resilience is defined as the distance between the stable and unstable state. Stability is defined as how quickly a dynamical system can return to its stable equilibrium after a perturbation. (B). The stability-resilience metrics quantified from effective potential curves are: a) potential width, b) potential depth, and c) potential slope. Potential width (a) directly quantifies the extent of perturbation a species could withstand before transitioning to another state (shown by the black ball in the effective potential curves), (b) potential depth, quantifies the resilience in the broad sense and higher value relates to higher resilience, and (c) potential slope in the figure quantifies stability of the population or how fast a system could bounce back following a perturbation. Dashed black lines in the community motifs represent predator-prey interactions, solid maroon lines represent mutualistic interactions, and solid blue lines represent competitive interactions. Vertical dashed blue line at t=200 denotes the timepoint at which environmental perturbation starts.

#### Community 1 - Mutualistic interactions

For a six species community with mutualistic interactions (e.g., a plant-pollinator), we define *r*’(*t*) as 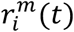, where growth rate of a species *i* depends on co-evolutionary interactions between the species in the community, and superscript *m* denotes mutualistic community. Specifically, we model mutualistic interactions using trait-matching functions with a linear functional response. The growth equation for a species engaging in mutualistic interactions can be written as:

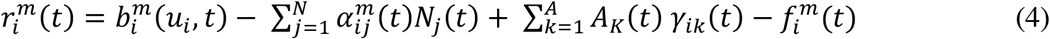

In this model, 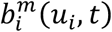 is the growth rate in the absence of competition and mutualism; *N_j_*(*t*) is the density of competitors; *A_K_*(*t*) is the density of the other group of species that provides the mutualistic benefits. We specifically ensure that eco-evolutionary dynamics are primarily dependent on trait-mediated mutualistic interactions, 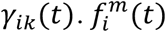 is the environmental forcing function that forces a population through a fold bifurcation. *γ_ik_* (*t*) can be written as (for details of the model and parameters used see supplementary 1):

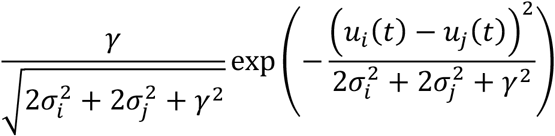

#### Community 2 – predator-prey interactions

We used a tri-trophic predator-prey model, with one top predator (species 6 in the predator-prey motif, Fig. 1), two consumers (species 4 and 5) and three basal species (species 1, 2 and 3; see Fig. 1). Predation on species 4 and 5 by the top predator (species 6) was possible through a Gaussian trait-matching function. Similarly, species 4 and 5 were able to predate on the basal species if their trait means were similar to the basal species (Haney and Siepielski 2018). Specifically, the growth rate of a basal species, 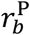 (superscript *P* denotes predator-prey community), in the predator-prey food web model depends on the inter- and interspecific competition, and trait-mediated predation from consumers (species 4 and 5) given as:

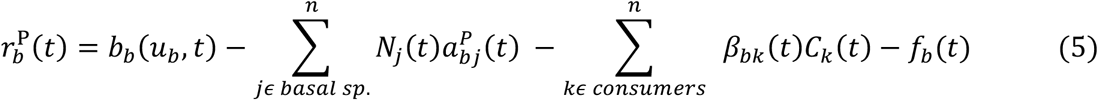

Here, *b_b_*(*u_b_*, *t*) is the per capita growth rate of a basal species in the absence of competition and predation, and dependent on the mean trait (*u_b_*). Note that the subscript *b* denotes the basal species and the superscript P denotes the community, which here is the predator-prey. Intra- 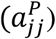 and interspecific 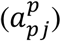 competition for the basal species are fixed (i.e., trait-independent) with intraspecific competition being substantially larger than interspecific competition. *β_bk_* (*t*) is the trait-mediated predation on a basal species *b* by the consumer species *k* which are species 4 and 5.

For the consumer species 4 and 5, the growth rate, 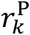, is defined as:

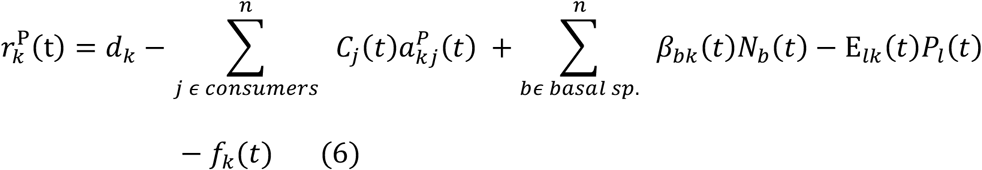

And for the top predator, growth rate, 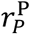, is defined as:

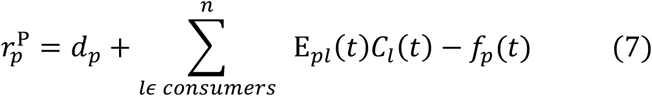

Here, 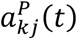 in equation 6 is the intra- and interspecific trait-independent competition among the consumer species (species 4 and species 5). Intraspecific competition in basal species was fixed at 0.5, whereas for the consumers were fixed at 0.1 (Petchey et al. 2008). Similarly, interspecific competition for basal species was drawn randomly from a uniform distribution [0, 0.05]; for the consumer species interspecific interactions were randomly drawn from a uniform distribution ranging from [0, 0.005], signifying weak interspecific competition at higher trophic levels (Petchey et al. 2008); *d_p_, d_k_* is the per capita mortality of top predator and the consumers respectively. We specifically followed (Petchey et al. 2008) to parameterize the mortality rate of the consumers (*d_k_*) and the top predators (*d_p_*). Since top predators or consumers at higher trophic levels have lower mortality rates than basal species (Cohen et al. 2003), we drew mortality rates from a uniform distribution [-0.02, −0.001] and sorted the mortality rates in a way that the top predator had the lowest mortality and the consumer species had a slightly higher mortality rate. *β_bk_* (*t*) in equation 6 is the trait-mediated predation on basal species which was a gaussian interaction kernel (Haney and Siepielski 2018); *E_lk_*(*t*) is trait-mediated predation by the top predator on the consumer species 4 and 5, also a gaussian function (see supplementary 2 for details of the formulation and parameters used in the model). Finally, *f_p_*(*t*), *f_b_*(*t*), *f_c_*(*t*) are the environmental forcing function for the top predator, the basal species and for the intermediate consumers respectively that forces a species though a fold bifurcation (Clements & Ozgul 2016).

### 2.2 Simulating population collapses

Before simulating species collapse experiments, we first assembled the communities. The starting point for assembling stable six-species communities was initial mean trait values for the six species for each of the community type. For both predator-prey community and the mutualistic community, we sampled and assigned mean trait values randomly from a uniform distribution in the range [-0.5, 0.5]. Based on these mean trait values, trait-based interactions either resulted in feasible communities or unfeasible communities. In addition, in all of our simulations for both the community types, we sampled competition coefficients in a way that intraspecific competition was always greater than interspecific competition. This specific rule ensured that trait-independent competitive interactions did not become the reason for unfeasible communities both in mutualism and predator-prey communities. Whenever simulations with certain random mean trait values, competitive coefficients, growth rates and death rates led to feasible communities we used those mean trait values and other parameters as a starting point for our next collapse experiments.

Once we had the mean trait values of all the six species, competitive coefficients, growth/death rates that resulted in feasible communities, we used these parameters to then simulate population collapses. Instead of perturbing the whole community over time (Dakos and Bascompte 2014), only one of the six species was perturbed until it collapsed, a scenario mirroring, for example, the overharvesting of a particular fish stock (Link and Watson 2019) or species specific disease (Castro and Bolker 2005). Forcing of each species in each of the community was done by fold stochastic harvesting after a period of time which allowed for transitory dynamics in the system (*t* = 200). Each species’ collapse experiment had 100 replicates, producing a total of 600 collapses for a community type (see fig. S2-S4 for example timeseries).

### 2.3 Effective potential curves

The resilience and stability for alternative stable states in complex systems can be visualized with the help of a ‘ball and cup’ diagram (also known as stability-landscape or effective potential curves), where the ball represents the state of the dynamical system and the cup would represent the effective potential energy (Beisner et al. 2003, Nolting and Abbott 2015) (see fig. 1). How stable and how resilient the population is to external perturbation depends not only on intrinsic ecological factors such as growth rate or plasticity to environmental change (Reed et al. 2010, Baruah et al. 2020), but also on the type of interactions it is engaging in (Barabás et al. 2017). A species embedded in either of the communities described above can be represented in the form of 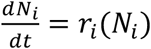, with the effective potential of the dynamical system then being:

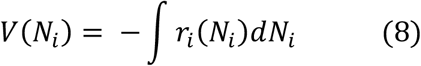

Both the different community types we can estimate *V*(*N_i_*) as a function of other species at any particular time *t*. For instance, for a mutualistic community, *V*(*N_i_*) at any time *t* could be written as (see supplementary 3 for *V*(*N_i_*) for the other community type):

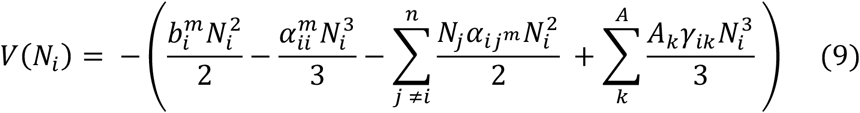

Where *N_j_*, is population density of species *j* at time *t* and 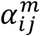 is interspecific effect of species *j* on species *i* at time *t*, and superscript *m* stands for mutualistic community; *a_ii_* is the intraspecific effect at time *t* and 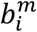 is the growth rate at time *t*. We can then quantify *V*(*N_i_*) for different values of and plot the potential curve at a particular time point *t*. Specifically, we estimate *V*(*N_i_*) just before environmental forcing starts i.e., at t = 200 time point (see Figure 1). At that time point, we estimated three quantities that accurately characterize the potential curves: a) scaled potential width, which is the distance between the alternative state and the lowest point of the potential. Scaled potential width can be quantified as population density at the lowest effective potential value subtracted by population density at the highest effective potential value which then was scaled by maximum population density (Nolting and Abbott 2015); b) depth of the potential curve given as: min(*V*(*N_i_*) - max(*V*(*N_i_*), which quantifies how resilient the system is, and c) the slope of the potential curve which quantifies how stable the one-dimensional system is. We quantify these three metrics prior to the start of environmental forcing for each replicate simulation of population collapses (n=100) and for each of the community motifs, and evaluate whether different interaction types modify these stability-resilience metrics and in turn influence the utility of EWS of population collapses (see below).

In addition to quantifying stability metrics from effective potential curves, we also estimated gains in growth for all the species based on mutualistic interactions for the mutualistic community. For the predator-prey food-web community, we estimated increases in growth rate for the basal species; increases in growth rate for the intermediate consumers from predation and increases in growth rate for the top predator from predation on intermediate consumers. This was possible as we tracked abundance, trait, and trait-mediated interactions. We thus evaluated whether increases in growth rate through mutualistic interactions, or predation before the start of environmental perturbation relate to the performance of EWS in forecasting population collapse.

#### Population-level signals

##### Abundance-based EWS

We evaluate whether abundance-based EWS estimated from the abundance of single species could be affected by interactions with other species. We estimated standard deviation (SD) and autocorrelation (AR1) (using the ‘*earlywarnings’* package (Dakos et al. 2012)) from the abundance of each of the six species for each of the communities from the start of the perturbation till the time point of 45% decline of the species from its initial starting density. The rest of the time series was discarded. The challenge is to detect EWS before a significant species decline (>50%), which was why we used 45% decline in focal species as a cut-off for our analysis. One could also use the entire timeseries (100% decline in focal species), but doing so negates the effectiveness of predicting future collapses with EWS as a negative trend in abundance would be good enough to predict collapses. We used gaussian detrending to remove any trend in the abundance data and used 50% of the time series as the rolling window size. The expectation was thus that abundance-based EWS such as SD or AR1 should increase and predict population collapse before a significant decline. To estimate the increase in SD or AR1 over time before population collapse, we quantified Kendall’s tau correlation coefficient over time. Kendall’s tau rank correlation coefficient has values ranging from −1 to 1, where 1 indicate perfect positive correlation and −1 indicates perfect negative correlation. High positive Kendall’s tau value would indicate strong signals of population collapse and negative values would indicate false negatives i.e., a species collapsed but EWS failed to predict it.

##### Trait-based EWS

Trait-based EWS were estimated from the mean trait dynamics of the focal species being forced to collapse. The length of the trait time series used for trait-based EWS was equal to that of its corresponding abundance time series. Shifts in the mean trait of the species before 45% population decline were considered to be an indicator of population collapse. As a species was forced to collapse through harvesting, mean trait of the focal species could either remain unaffected or shift in response to increases in harvest pressure. Such a trait shift could be in any direction i.e., mean trait value could either increase or decrease. We thus considered shift in mean trait value in any direction as a potential warning signal. However, for the sake of easier analysis, whenever a shift in mean trait value was negative, we multiplied the mean trait of a species by −1 such that shift in mean trait value was always positive. Doing so ensured that an increase in the mean trait value would mean a positive Kendall’s tau value which we used to quantify strength of EWS in general.

#### Community-level signals

##### Abundance- and trait-based warning signals

In addition to the population level analyses described above, we also assessed whether measures of stability were detectable in community level data that comprised of two time series: (1) the sum of all of the abundances of all the species in the community at each time step, and (2) the mean trait value of all the species in the community at each time step. Thus, community-based indicators comprised of trends in *community SD, community AR1* (Strahan et al. 2016), and *community trait.* All community level measures were calculated and their change measured in the same way as the species level signals (see above). As with the species level analyses, community level signals were analyzed from the start of the environmental perturbation to the time point of 45% decline of the focal species that was being continuously perturbed. Trends in these EWS were measured using Kendall’s tau correlation coefficient, where positive values indicate positive strength in forecasting collapse.

##### Magnitude and direction of community variance

In addition to the community level warning signals analyses described above, we also consider an alternative metric of community stability. Multispecies communities are high-dimensional in nature with many state variables (e.g., species abundances). As such, when a high-dimensional system is perturbed continuously, fluctuations of certain state variables become much larger and slower than others as it approaches a transition, which can be captured by calculating the variance-covariance matrix. One of the main properties of such a covariance matrix is that its dominant eigenvector would represent the direction of the largest variance in the community and the dominant eigenvalue would represent the magnitude of this variance. As a consequence, changes in the dominant eigenvalue estimated from the covariance matrix would thus indicate a system’s sensitivity to external perturbation (Dakos 2017). We thus propose the ‘dominant eigenvalue’ as another class of warning signal of a community’s sensitivity to external perturbation. A similar class of warning signals were defined for spatially heterogenous systems (Chen et al. 2019) and for a competitive community via loadings derived from eigenvector decomposition (Dakos 2017). Such a class of indicator derived from the covariance matrix could only be applicable to systems that are high-dimensional in nature. The variance-covariance matrix C can be estimated from multispecies abundance time series with a rolling window of size *n* as

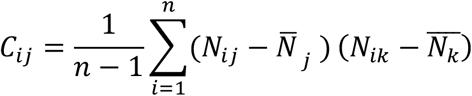

where N_ij_ is the abundance for species *i* for the *i_th_* time point, *n* is the size of the rolling window, 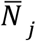 is the mean abundance over the rolling window time, *N_ik_* is the abundance of species *k* for the *ith* time point. Dominant eigenvalue of this covariance matrix calculated at each time point of the rolling window is the final class of warning signal used in this study. A shift in this dominant eigenvalue would thus infer increasing instability of the community. As with the other metrics the trend in the dominant eigenvalue through time was measured using Kendall’s tau correlation, facilitating direct comparison between all of the measures of stability.

## Results

### Mutualistic community

Measures of stability and resilience metrics from effective potential curves indicated that species 5 and 6 had higher potential slope and depth indicating higher stability and resilience among all the other species (Fig 3A-C). In addition, species 5 and species 6 had the strongest mutualistic benefits (Fig. 4A). Consequently, species-level AR1 and SD was able to forecast collapse of species 5 and species 6 better than all the other species (Fig. 2A).

**Figure 2.**
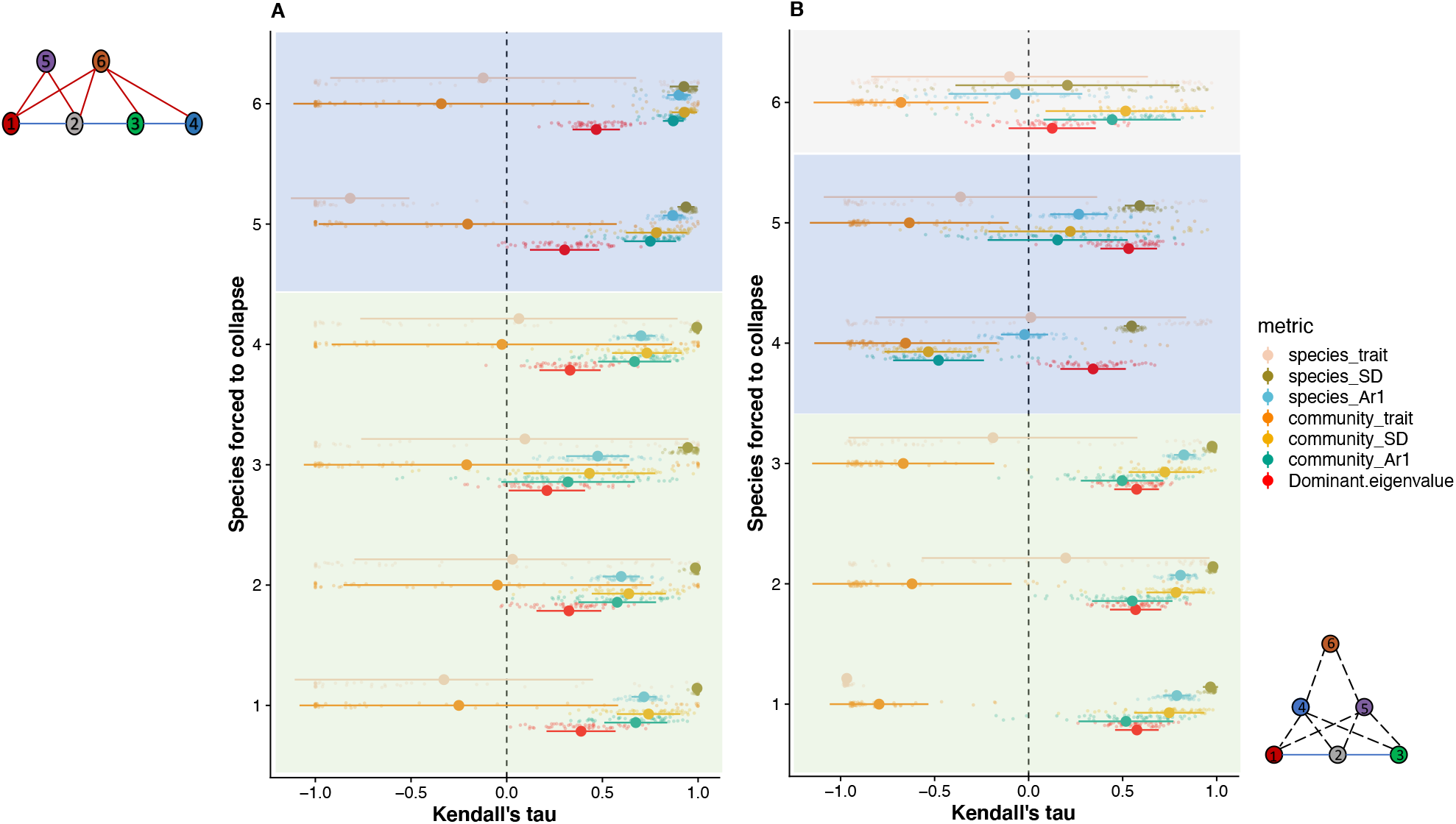
Mean Kendall’s tau value with standard error bars of community-based, and traitbased EWS of 100 replicate simulations of population collapse for mutualism only community (A), and predator-prey community (B). The color blocks highlight different trophic levels. From (A): For SD and ARI, measured at both species and community levels, Kendall’s tau was highest when species 5 or 6 (trophic level 2) were the target of forcing. From (B): EWS of population collapses for predator-prey community.

**Figure 3:**
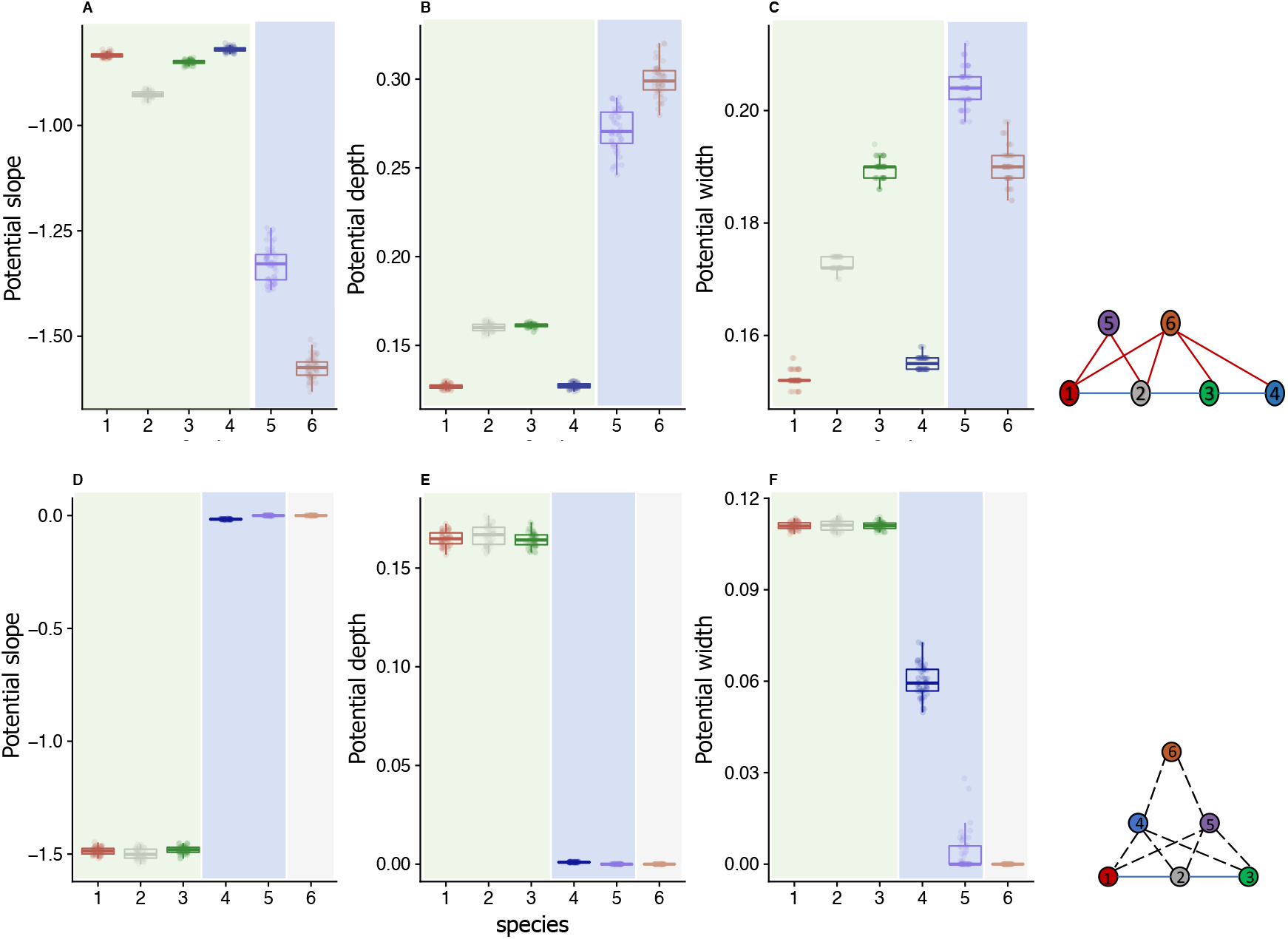
Metrics estimated from effective potential curves for mutualism-only community (A, B, C) and predator-prey community (D, E, F). The color blocks indicate different trophic levels (two for mutualism community and three for predator-prey community). Note that different shading areas represent different trophic levels. From A, B, C: Species 5 and 6 had the highest potential depth and steepest potential slope in comparison with all the other species for the mutualism-only community. Species 5 and 6 also had higher potential width when compared to others. As a consequence, species 5 and 6 showed the strongest community and abundance-based EWS (Fig. 2A). From D, E, F: all the basal species (species 1, 2, and 3) had higher potential width in comparison with the consumer (species 4 and 5) and the top predator species (species 6). In terms of potential slope (D) and depth (E) basal species had the highest among all the species indicating high stability which resulted in stronger EWS than others (Fig. 2B).

**Figure 4.**
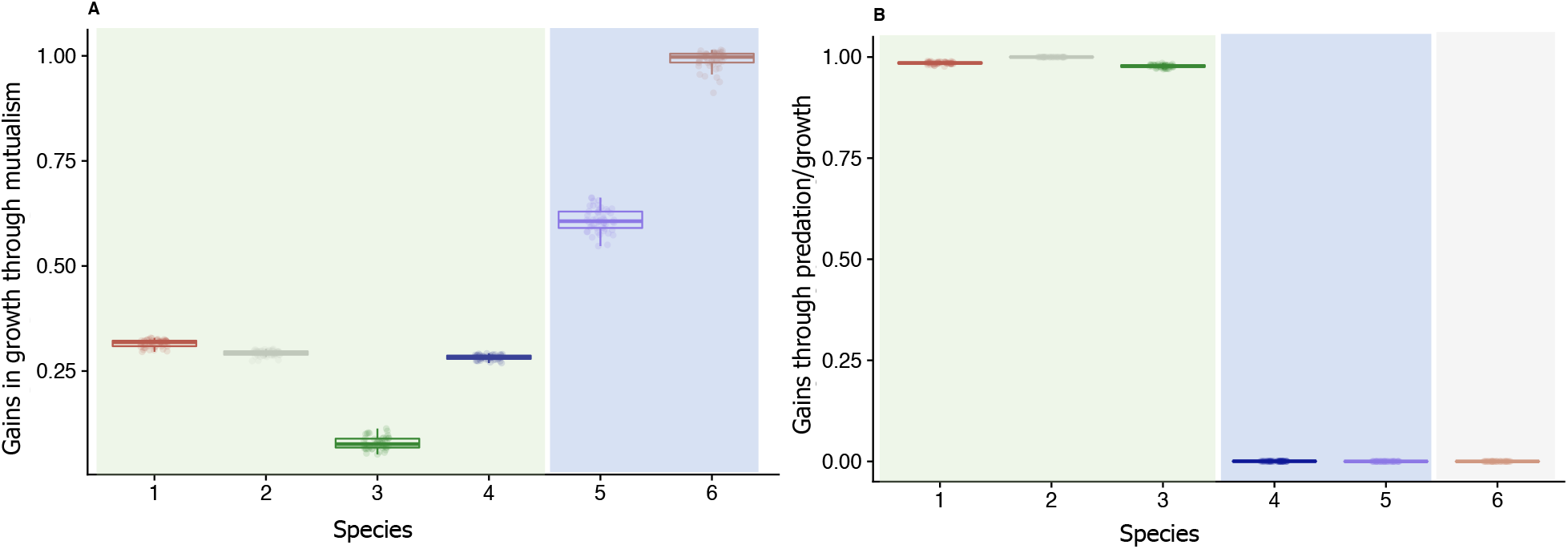
A) Gains in growth through mutualistic interactions, (B) predation for consumers and top predators, or density-independent growth of basal species. Shaded regions highlight the trophic levels in mutualism-only (A) and predator-prey community (B). In (A) species 5 and 6 had the highest mutualistic growth benefits. In (B) the basal species had the highest benefits in growth in comparison to the consumers and the top predator.

Mean trait-based EWS measured at the species level was an unreliable predictor of species collapse indicating that shifts in mean traits were not observed despite drastic declines in population sizes (Fig. 2A). In addition, community level trait-based indicators were also not significantly better in forecasting species level collapse compared to the abundance-based EWS. Dominant eigenvalue performed relatively well in forecasting species level collapse.

### Predator-prey community

The effective potential curves estimated prior to the onset of environmental perturbation indicated that species in the lowest trophic level (species 1-3) had the steepest slopes and highest depths and potential widths, in comparison to intermediate consumers (species 4 and 5) and top predator (species 6; Fig. 3D-F). This in turn led to a similar pattern in the strength of abundance-based EWS, with species-level AR1 and species-level SD displaying strong temporal trends in the basal species and weaker trends in the predatory species (Fig. 2B). Community level abundance-based EWS such as community AR1 and community SD performed relatively better in forecasting the collapse of basal species (species 1, 2, and species 3) than they did in forecasting collapse of predators (Fig. 2B). Dominant eigenvalue also performed similarly to abundance-based EWS in terms of forecasting species collapses (Fig. 2B).

In contrast, trait-based EWS whether it was measured at the community level or at the species level were unreliable and performed poorly in forecasting species collapse (Fig. 2B).

## Discussion

Whilst much work has been done in seeking EWS in simulated, experimental, and wild population data, thus far the potential of species interactions to influence the presence and reliability of these signals has been largely overlooked (Dakos 2017). We model eco-evolutionary dynamics of species in two simple but commonly considered community motifs (Cortez 2016, McPeek 2017). In our models, interactions between species are dynamic and trait-mediated, such that feedback loops can occur between changes in the ecological dynamics and the mean trait values of each species. Such feedbacks in our models allowed us to explore signals of collapse at different levels of biological organization, from trends in species traits to variance in community-level abundance data. Here, we evaluate the utility of phenomenological warning signals in forecasting impending collapses of species residing in different community motifs and demonstrate that the role of a species in a community can have significant impacts on our ability to forecast its loss. Whilst our network motifs are clearly simplifications of potential community structure observed in nature, we use these motifs to demonstrate the potential importance of interaction type in determining our ability to forecast species loss.

In the simulated mutualistic communities, species which had the strongest mutualistic interactions (i.e., species 5 and species 6) that led to overall high gains in growth (Fig 4A) had the deepest potential curve and the steepest slope among all the other species, and were thus the most stable. This strong mutualistic interaction and deeper potential curves meant that abundance-based EWS measured at the species level i.e., SD and AR1, performed significantly better in forecasting the collapse in these two species when compared to others in the mutualistic network.

In our simulated predator-prey communities, the strength of species level abundance-based EWS (AR1 and SD) in forecasting population collapse were most prominent in the basal species in comparison to when intermediate consumers or the top predator were forced to collapse (Fig 2B). Specifically, the basal species had a steeper and a deeper potential curve indicating higher stability and resilience in comparison to the intermediate and top predators (Fig 3 D-F). Interestingly, these basal species also had higher growth rate in comparison to the consumers and top predators (Fig. 4B), which in turn reflected both in the metrics estimated from the potential-well curves and on the performance of the abundance-based EWS (Fig 3D-F, 2B).

AR1 and SD, measured at the species level and at the community level, were not reliable in forecasting species collapse when species 4, 5 (which predated on the basal species), and species 6 (which was the top predator) were forced to collapse. This was also due to the fact that these three species had the shallowest potential curve among all the species before environmental perturbation started and had the least fitness benefits from predation (Fig 3D, Fig. 4B).

Abundance-based community level indicators, particularly community SD and community AR1, were reliable in forecasting species level collapses when species 5 and 6 were forced to collapse in the mutualistic network. Such a result strongly indicated that stronger mutualistic interactions that positively influences growth rate (Fig. 4A) could contribute more to community level dynamics and stability as opposed to weaker mutualistic interactions that could lead to weaker gains in growth as observed in species 1, 3 and 4. In our mutualistic network, species 5 and 6 were benefitting from four mutualistic interactions with the four other species thereby leading to higher gains in growth in comparison to species 1, 2,3 and 4, which only gained mutualistic benefits from the two species (species 5 and 6). When any of the four species (1, 2, 3 or 4, see figure 1 mutualism-only motif) were forced to collapse, the loss in mutualistic benefits for species 5 and 6 were not large enough to impact community wide stability. For instance, if species 1 was forced to collapse, species 5 and 6 could still maintain positive fitness benefits because of their mutualistic interaction with the remaining three species. Hence, decline in species 1 numbers due to forcing did not influence abundances of species 5 and 6. Thus, when any of the species 1, 2, 3, or 4 was forced to collapse, decline in their numbers did not impact community dynamics and community-wide signals were not strong enough to indicate community-wide instability. On the contrary, when species 5 or 6 were forced to collapse, the loss in fitness for the other four species (species 1,2,3, and 4) were much larger as they had fitness benefits *only* from mutualistic interactions with species 5 and 6, and loss of one of these two species had a significant impact on community dynamics. In turn, community-based signals performed much better when species 5 or 6 were forced to collapse in comparison to others. This also indirectly indicated that the degree distribution of the mutualistic network could impact community stability. Previous studies have suggested that properties of network architecture such as connectance, degree distribution, or nestedness could impact stability and robustness of communities to species loss (Bastolla et al. 2009, Dunne and Williams 2009). Our two community modules varied in terms of connectance and trophic levels. Connectance and nestedness, for instance, could increase the likelihood of occurrence of abrupt collapses in mutualistic networks (Lever et al. 2014, Baruah 2022). Similarly, higher food-web connectance has been linked with community instability (Landi et al. 2018). Thus, it is possible that such network properties could influence which species could exhibit stronger signals than others, however, a detailed further investigation is required.

In the predator-prey communities, abundance-based community-level indicators (community AR1, community SD) were also reliable in forecasting species level collapse, particularly for the basal species (Fig. 2B). Such a result indicated that the basal species had the strongest influence on community stability. Decline in the basal species numbers had to have an impact on the overall community dynamics for the occurrence of any observable increase in community level indicators. In addition, the predator species were predominantly dependent on the basal species for growth, and as the basal species numbers declined due to environmental forcing, the predator species, consequently, were negatively impacted (Petchey et al. 2008). Thus, there was a rise in the community level indicators over time indicating community instability (Fig. 3A). On the other hand, when the predator species were forced to collapse, community-level indicators were unreliable in forecasting species level collapses, exhibiting a high number of false negatives (the error bars being large and crossing the zero line in Fig. 2B). Thus, our simulation results clearly indicate that predatory species were the most vulnerable to environmental perturbation and were liable to collapse without exhibiting strong signals of population collapse.

Species level trait-based signals in our models included mean trait measured at the species level. Contrary to previous studies where it was consistently shown that trait-based signals outperformed abundance-based EWS (Clements and Ozgul 2016, Baruah et al. 2019, 2020, Arkilanian et al. 2020), our study on multispecies communities consistently suggested that trait-based indicators were less reliable irrespective of whether a species was in a mutualistic or in a prey-predator community (Fig 2A-B). This unexpected result may be derived from the way in which mean phenotypic traits influenced population dynamics. Our modelling approach forced populations to collapse through harvesting of a species that then goes through a fold bifurcation. This directly impacted population dynamics, but only influenced mean phenotypic dynamics indirectly. Thus, unless there was a drastic change in population size, the impacts on mean phenotypic dynamic would be relatively small. In addition to this, if however, an environmental perturbation impacted phenotypic traits directly, than the potential for traits to be used as an indicator increases significantly (Baruah et al. 2019). Such instances could be when there was a decline in food availability that could impact phenotypic traits such as body size, or a size-based harvesting regime that could also lead to a shift in body size (Clements and Ozgul 2016, Burant et al. 2021). However, our environmental perturbation did not have direct impacts on mean traits directly, and hence observable change in mean phenotypic dynamics were negligible, leading to poor performance of trait-based signals measured at the species level. Similar to species-level trait-based EWS, community-level trait-based EWS were unreliable in forecasting species level collapse regardless of which community type the species belonged. Since declines in species abundances resulting from environmental forcing did not impact species mean traits directly, shifts in both mean trait value or community trait value were not observed.

Our study also evaluated another indicator that took into account the variance and covariances of abundance of multiple species over time (which we call the dominant eigenvalue), similar to the one derived from eigenvector decomposition of a high-dimensional competitive system (Dakos 2017). A similar approach has recently been tested in predicting transitions in spatial connected populations (Chen et al. 2019). Rise in dominant eigenvalue was most prominent in collapse of species which had the steepest and the deepest potential curve, and was particularly effective in forecasting collapse of species in the predator-prey food-web community. It is to be noted that the dominant eigenvalue estimated from the variance-covariance matrix is not an indicator measured at the species level. In comparison to all the other indicators, dominant eigenvalue performed fairly well in forecasting species-level collapse. Particularly, in some cases when community level and species level indicators were underperforming, for instance for species 4 and 5 (intermediate predators) in the predator-prey community, dominant eigenvalue outperformed community-level (community Ar1, community SD) and species-level indicators (species Ar1 and SD) (Fig. 2B).

Our modelling exercise ignored the spatial extent of ecological systems. Recent study had addressed that self-organization or spatial pattern formation could be linked to evasion of tipping points instead of signaling instability of spatial ecosystems (Rietkerk et al. 2021). One avenue of future research would be to extend our framework to include spatial system and understand how spatial instability could be linked to local community structure. Also, to be noted, that our approach could be only useful in systems that have a potential. Dynamical ecological systems such as predator-prey (Cortez 2016), consumer-resource (Holt and Polis 1997) or mutualistic systems (Dakos and Bascompte 2014, Baruah 2022), or lake-eutrophication models (Carpenter et al. 2011, Nolting and Abbott 2015) are gradient systems and as such should display stability-landscape curves. However, our approach of combining metrics from stability-landscape curve with phenomenological EWS would not work for non-gradient ecological systems or systems devoid of any effective potential.

In conclusion, our results demonstrate that the structure of the community in which a species resides can fundamentally change how predictable the collapse of that species’ population is. Whilst community-level abundance-based warning signals often provided the most reliable predictions of population collapse; in reality such time series are difficult to collect and rarely available. However, for systems such as fisheries where the strength of interactions is known and community level times series data are often collected, our results suggest dominant eigenvalue estimated from the variance-covariance matrix could potentially be useful in forecasting species collapses. However, in the absence of such community level data, we highlight the potential of using indicator species as reliable forecasters of the collapse of species and communities. Such indicator species may be identifiable by identifying where the strongest interaction strengths in a community are, and then measuring these species for signals of instability. Critically, that the type of community a species resides in may influence how predictable collapse is raises concerns about the generality of EWS frameworks, and their utility in predicting real-world collapse.

## Acknowledgements

The authors declare no conflicts of interests. The study was funded by Forschungskredit UZH FK-18-082 to GB, ERC-FP7 #337785 and SNSF 310003A_146445 to AO.

## Data availability

The data and R scripts will be uploaded to Zenodo.

## References

Arani, B. M. S., S. R. Carpenter, L. Lahti, E. H. van Nes, and M. Scheffer. 2021. Exit time as a measure of ecological resilience. Science.

Arkilanian, A. A., C. F. Clements, A. Ozgul, and G. Baruah. 2020. Effect of time series length and resolution on abundance- and trait-based early warning signals of population declines. Ecology 101:e03040.

Barabás, G., M. J Michalska-Smith, and S. Allesina. 2016. The Effect of Intra- and Interspecific Competition on Coexistence in Multispecies Communities. The American naturalist 188:E1–E12.

Barabás, G., M. J. Michalska-Smith, and S. Allesina. 2017. Self-regulation and the stability of large ecological networks. Nature Ecology & Evolution 1:1870–1875.

Baruah, G. 2022. The impact of individual variation on abrupt collapses in mutualistic networks. Ecology Letters 25:26–37.

Baruah, G., C. F. Clements, F. Guillaume, and A. Ozgul. 2019. When Do Shifts in Trait Dynamics Precede Population Declines? The American Naturalist 193:633–644.

Baruah, G., C. F. Clements, and A. Ozgul. 2020. Eco-evolutionary processes underlying early warning signals of population declines. Journal of Animal Ecology 89:436–448.

Baruah, G., C. F. Clements, and A. Ozgul. 2021. Effect of habitat quality and phenotypic variation on abundance- and trait-based early warning signals of population collapses. Oikos 130:850–862.

Bastolla, U., M. A. Fortuna, A. Pascual-García, A. Ferrera, B. Luque, and J. Bascompte. 2009. The architecture of mutualistic networks minimizes competition and increases biodiversity. Nature 458:1018–1020.

Beisner, B., D. Haydon, and K. Cuddington. 2003. Alternative stable states in ecology. Frontiers in Ecology and the Environment 1:376–382.

Boerlijst, M. C., T. Oudman, and A. M. de Roos. 2013. Catastrophic Collapse Can Occur without Early Warning: Examples of Silent Catastrophes in Structured Ecological Models. PLoS ONE 8.

Burant, J. B., G. S. Betini, and D. R. Norris. 2019. Simple signals indicate which period of the annual cycle drives declines in seasonal populations. Ecology Letters 22:2141–2150.

Burant, J. B., C. Park, G. S. Betini, and D. R. Norris. 2021. Early warning indicators of population collapse in a seasonal environment. Journal of Animal Ecology 90:1538–1549.

Carpenter, S. R., J. J. Cole, M. L. Pace, R. Batt, W. A. Brock, T. Cline, J. Coloso, J. R. Hodgson, J. F. Kitchell, D. A. Seekell, L. Smith, and B. Weidel. 2011. Early warnings of regime shifts: a whole-ecosystem experiment. Science (New York, N.Y.) 332:1079–82.

Castro, F. D., and B. Bolker. 2005. Mechanisms of disease-induced extinction. Ecology Letters 8:117–126.

Chen, S., E. B. O’Dea, J. M. Drake, and B. I. Epureanu. 2019. Eigenvalues of the covariance matrix as early warning signals for critical transitions in ecological systems. Scientific Reports 9:1–14.

Clements, C. F., J. L. Blanchard, K. L. Nash, M. A. Hindell, and A. Ozgul. 2017. Body size shifts and early warning signals precede the historic collapse of whale stocks. Nature Ecology & Evolution 1:1–6.

Clements, C. F., and A. Ozgul. 2016. Including trait-based early warning signals helps predict population collapse. Nature Communications 7:1–8.

Clements, J. M. Drake, I. G. Jason, and A. Ozgul. 2015. Factors Influencing the Detectability of Early Warning Signals of Population Collapse. The American Naturalist 186:50–58.

Clements, and A. Ozgul. 2018. Indicators of transitions in biological systems. Ecology Letters 21:905–919.

Cohen, J. E., T. Jonsson, and S. R. Carpenter. 2003. Ecological community description using the food web, species abundance, and body size. Proceedings of the National Academy of Sciences 100:1781–1786.

Cortez, M. H. 2016. How the Magnitude of Prey Genetic Variation Alters Predator-Prey Eco-Evolutionary Dynamics. The American naturalist 188:329–41.

Dai, L., D. Vorselen, K. S. Korolev, and J. Gore. 2012. Generic Indicators for Loss of Resilience Before a Tipping Point Leading to Population Collapse. Science 336:1175–1177.

Dakos, V. 2017. Identifying best-indicator species for abrupt transitions in multispecies communities. Ecological Indicators.

Dakos, V., and J. Bascompte. 2014. Critical slowing down as early warning for the onset of collapse in mutualistic communities. Proceedings of the National Academy of Sciences 111:17546–17551.

Dakos, V., S. R. Carpenter, W. A. Brock, A. M. Ellison, V. Guttal, A. R. Ives, S. Kéfi, V. Livina, D. A. Seekell, E. H. van Nes, and M. Scheffer. 2012. Methods for Detecting Early Warnings of Critical Transitions in Time Series Illustrated Using Simulated Ecological Data. PLOS ONE 7:e41010.

Dakos, V., S. R. Carpenter, E. H. van Nes, and M. Scheffer. 2014. Resilience indicators: prospects and limitations for early warnings of regime shifts. Philosophical Transactions of the Royal Society B: Biological Sciences 370:20130263–20130263.

Dakos, V., B. Matthews, A. P. Hendry, J. Levine, N. Loeuille, J. Norberg, P. Nosil, M. Scheffer, and L. De Meester. 2019. Ecosystem tipping points in an evolving world. Nature Ecology & Evolution 3:355–362.

Doncaster, C. P., V. Alonso Chávez, C. Viguier, R. Wang, E. Zhang, X. Dong, J. A. Dearing, P. G. Langdon, and J. G. Dyke. 2016. Early warning of critical transitions in biodiversity from compositional disorder. Ecology 97:3079–3090.

Drake, J. M., and B. D. Griffen. 2010. Early warning signals of extinction in deteriorating environments. Nature 467:456–9.

Dunne, J. A., and R. J. Williams. 2009. Cascading extinctions and community collapse in model food webs. Philosophical Transactions of the Royal Society B: Biological Sciences 364:1711–1723.

Dutta, P. S., Y. Sharma, and K. C. Abbott. 2018. Robustness of early warning signals for catastrophic and non-catastrophic transitions. Oikos.

Guimarães, P. R., V. Rico-Gray, P. S. Oliveira, T. J. Izzo, S. F. dos Reis, and J. N. Thompson. 2007. Interaction Intimacy Affects Structure and Coevolutionary Dynamics in Mutualistic Networks. Current Biology 17:1797–1803.

Haney, S. D., and A. M. Siepielski. 2018. Tipping Points in Resource Abundance Drive Irreversible Changes in Community Structure. The American Naturalist 191:668–675.

Hastings, A., K. C. Abbott, K. Cuddington, T. Francis, G. Gellner, Y.-C. Lai, A. Morozov, S. Petrovskii, K. Scranton, and M. L. Zeeman. 2018. Transient phenomena in ecology. Science (New York, N.Y.) 361:eaat6412–eaat6412.

Holt, R. D., and G. A. Polis. 1997. A Theoretical Framework for Intraguild Predation. The American Naturalist 149:745–764.

Kéfi, S., V. Dakos, M. Scheffer, E. H. Van Nes, and M. Rietkerk. 2013. Early warning signals also precede non-catastrophic transitions. Oikos 122:641–648.

Lade, S. J., and T. Gross. 2012. Early warning signals for critical transitions: A generalized modeling approach. PLoS Computational Biology.

Landi, P., H. O. Minoarivelo, Å. Brännström, C. Hui, and U. Dieckmann. 2018. Complexity and stability of ecological networks: a review of the theory. Population Ecology 60:319–345.

Lever, J. J., I. A. van de Leemput, E. Weinans, R. Quax, V. Dakos, E. H. van Nes, J. Bascompte, and M. Scheffer. 2020. Foreseeing the future of mutualistic communities beyond collapse. Ecology Letters 23:2–15.

Lever, J. J., E. H. van Nes, M. Scheffer, and J. Bascompte. 2014. The sudden collapse of pollinator communities. Ecology Letters.

Link, J. S., and R. A. Watson. 2019. Global ecosystem overfishing: Clear delineation within real limits to production. Science Advances 5:eaav0474.

McPeek, M. A. 2017. Evolutionary community ecology.

Mougi, a., and M. Kondoh. 2012. Diversity of Interaction Types and Ecological Community Stability. Science 337:349–351.

Nolting, B. C., and K. C. Abbott. 2015. Balls, cups, and quasi-potentials: quantifying stability in stochastic systems. Ecology 97:15–1047.1.

Petchey, O. L., A. Eklöf, C. Borrvall, and B. Ebenman. 2008. Trophically Unique Species Are Vulnerable to Cascading Extinction. The American Naturalist 171:568–579.

Rankin, D. J., and A. López-Sepulcre. 2005. Can adaptation lead to extinction? Oikos 111:616–619.

Reed, T. E., R. S. Waples, D. E. Schindler, J. J. Hard, and M. T. Kinnison. 2010. Phenotypic plasticity and population viability: the importance of environmental predictability. Proceedings. Biological sciences / The Royal Society 277:3391–400.

Rietkerk, M., R. Bastiaansen, S. Banerjee, J. van de Koppel, M. Baudena, and A. Doelman. 2021. Evasion of tipping in complex systems through spatial pattern formation. Science.

Scheffer, Bascompte, W. A. Brock, V. Brovkin, S. R. Carpenter, V. Dakos, H. Held, E. H. van Nes, M. Rietkerk, and G. Sugihara. 2009. Early-warning signals for critical transitions. Nature 461:53–59.

Spanbauer, T. L., C. R. Allen, D. G. Angeler, T. Eason, S. C. Fritz, A. S. Garmestani, K. L. Nash, J. R. Stone, C. A. Stow, and S. M. Sundstrom. 2016. Body size distributions signal a regime shift in a lake ecosystem. Proceedings of the Royal Society B 283:20160249–20160249.

Strahan, R. T., A. J. S. Meador, D. W. Huffman, and D. C. Laughlin. 2016. Shifts in community-level traits and functional diversity in a mixed conifer forest: a legacy of land-use change. Journal of Applied Ecology 53:1755–1765.

Suweis, S., and P. D’Odorico. 2014. Early Warning Signs in Social-Ecological Networks. PLOS ONE 9:e101851.

