## Appendix_file for "Community structure determines the predictability of population collapse"

### 1. Mutualistic interaction

For the mutualistic interaction community, we assume that competition is trait-independent but mutualistic interaction is trait-dependent. Again, for mutualistic interactions, individual of trait $z$ of species $i$ (say plants) will interact strongly and have fitness gains from another species $j$ (say animals) in the community if their traits are similar. Hence one could write the per-capita growth rate for a species in either of the two trophic levels of mutualism community as:

$$r'_m(z,t,N) = \int \left( b_i^m(z) - \sum_{j=1}^{S} N_j(t)\alpha_{ij}^m(t) + \sum_{k=1}^{A} A_k(t) \int g(z,z')p_k(z',t)(t) \right) p_i(z,t)dz \tag{1}$$

Where, S is the number of species in the basal level (species 1, 2, 3, 4) and A is the number of species in other trophic level (species 5, 6).

where, $b_i^m(z)$ is a rectangular fitness function given as :

$$b_i^m(z) = \begin{cases} 1, & \text{if } \theta \geq z \geq -\theta \\ 0, & \text{otherwise,} \end{cases}$$

Thus, population dynamics of a species $i$ in either of the two trophic levels in the mutualism only community could be written as:

$$\frac{dN_i}{dt} = N_i \int \left( b_i^m(z) - \sum_{j=1}^{S} N_j(t)\alpha_{ij}^m(t) + \sum_{k=1}^{A} A_k(t) \int g(z,z')p_k(z',t)(t) \right) p_i(z,t)dz \tag{2}$$

$$= N_i \left( b_i^m(u_i) - \sum_{j=1}^{S} N_j(t)\alpha_{ij}^m(t) + \sum_{k=1}^{A} A_k(t)\gamma_{ik}(u_i,u_j) \right) \tag{3}$$

$\gamma_{ik}(u_i,u_j)$ is the average mutualistic interaction between two species $i$ (belonging to species 1,2 ) and species $j$ (belonging to 3,4,5,6) with trait means $u_i$ and $u_j$. The mean growth rate of a species in the trait axis i.e., $b_i^m(z)$ could be written as from above :

$$\int b_i^m(z)p_i(z,t)dz = \frac{1}{2}\left[ erf(\frac{\theta - \mu_i}{\sqrt{2}\sigma_i}) + erf(\frac{\theta + \mu_i}{\sqrt{2}\sigma_i}) \right]$$

where $\theta$ goes from -1 to 1 defining the boundary of the trait axis. Within this trait boundary, species will have a growth rate of 1 in the absence of mutualistic and competitive interaction.

Here, $A_k$ is the density of the mutualistic counterparts. In the mutualistic community module (fig. 1 main-text) , $A_k$ could represent either species 1 or 2 or species 3,4,5,or 6. If, for instance, $N_i$ represents any one of the two basal species (1 or 2), then $A_k$ would be the density of any species $k$ belonging to the other trophic level either species 3,4,5, or 6.

Intraspecific competition for all the species was kept at 0.5 and interspecific competition was randomly assigned during each simulation from a random uniform distribution $[0.001, 0.005]$ . Species density at the start of the simulation was kept at 1. Initial mean trait values were randomly assigned to each species until we found a set of mean trait values $u_i$ that led to feasible species densities. We used these mean trait values to start the simulations for species collapses. Species' trait variances for this community was randomly assigned for each

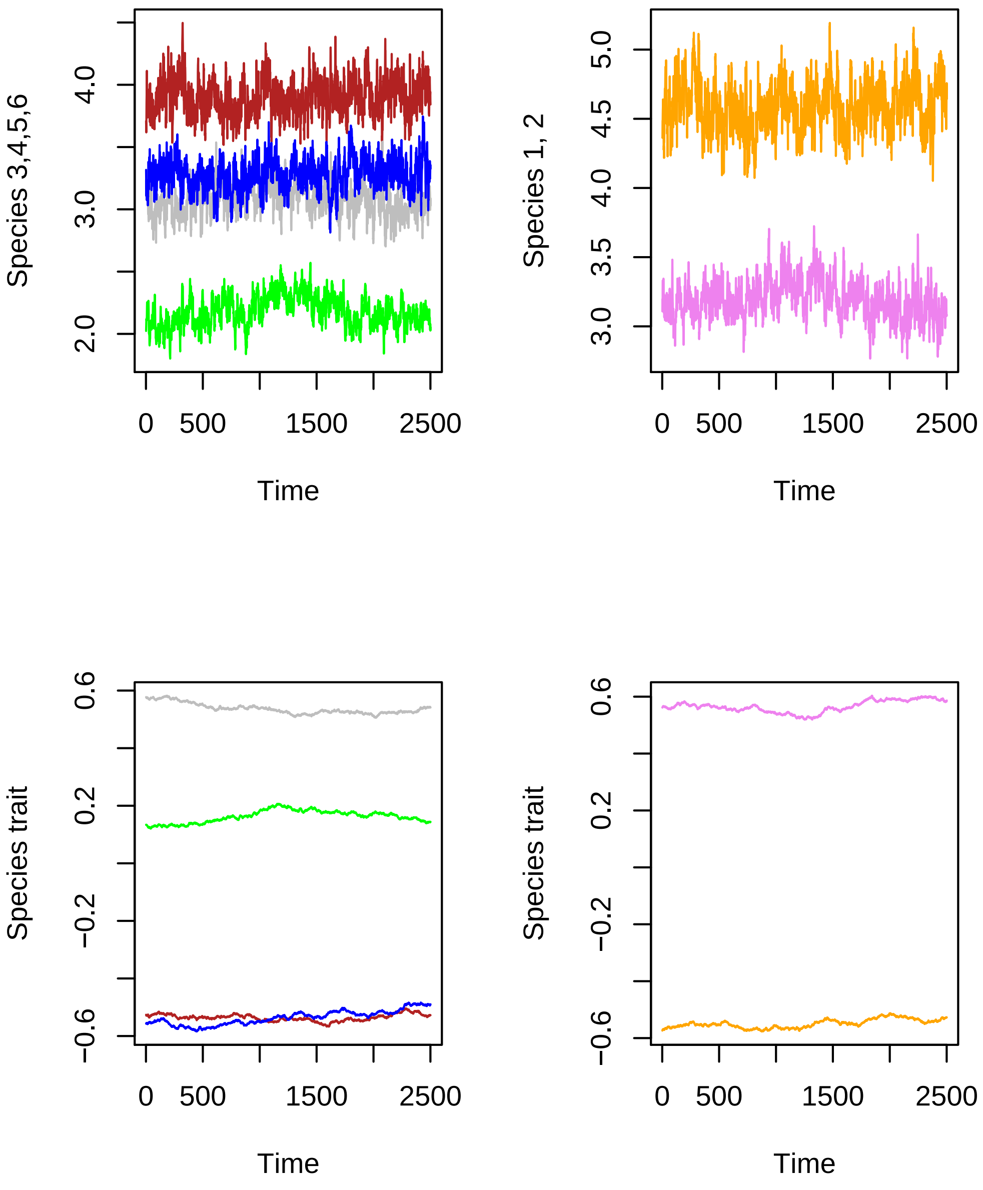


Figure S1: Time series of all six species with the parameter values mentioned in the appendix for the mutualistic interaction community.

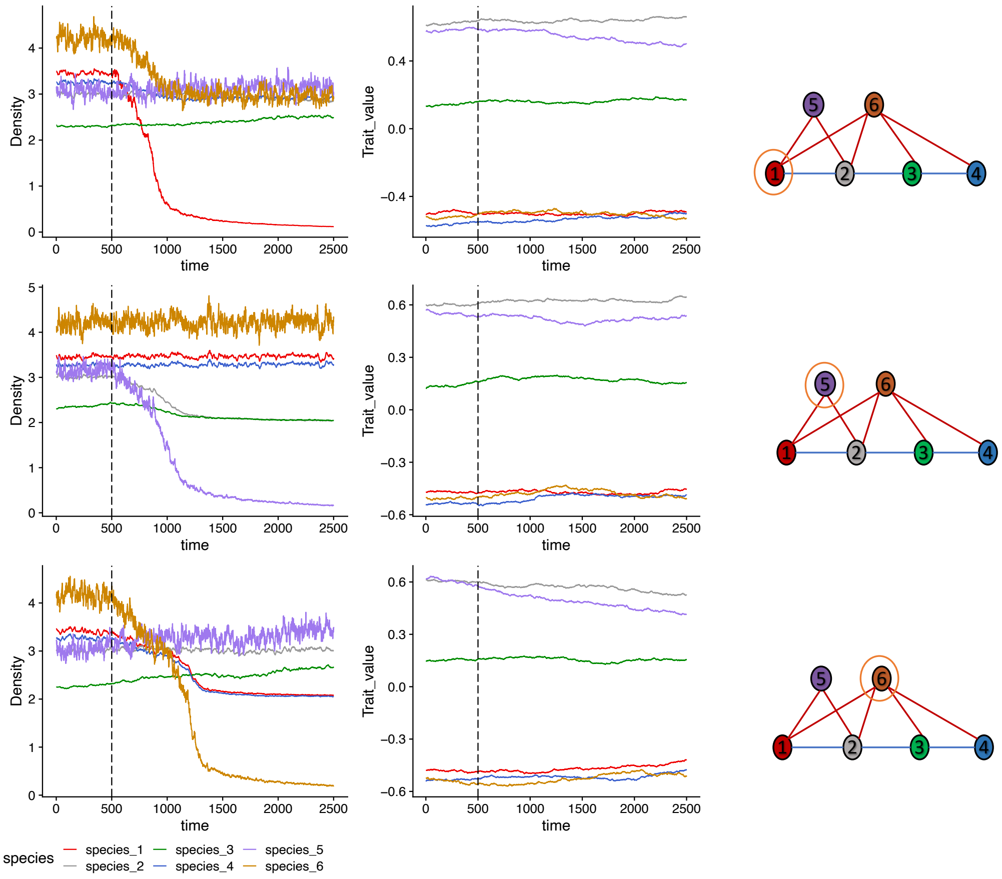


Figure S2: Time series of abundance and trait of all six species with the parameter values mentioned in the appendix for the mutualistic community. Dashed vertical line indicates the timepoint of species forcing. In the community motif circled species index number is the one being forced to collapse while rest of the species in the community remains unperturbed.

replicate simulation of population collapse from a random uniform distribution $[0.005, 0.02]$. At each time step of population dynamics we added Gaussian white noise with mean zero and variance of 0.5.

Mean trait dynamics $u_i$ of a species $i$ followed from equation 2 of the main-text.

Example time series of species collapse in mutualistic community are shown in figure S2.

### 2. Predator-prey interactions

In the community of predator-prey interactions, competion was trait-independent but predator-prey interactions were trait-mediated. Growth rate of species $i$ for the basal species can be written as:

$$r_p^P(z,t,N) = \int \left( b_p - \sum_{j\epsilon \ basal \ sp.} \alpha_{\mathrm{pj}}^P(t)\, N_j(t) - \sum_{k\epsilon \ consumers} C_k(t) \int b(z,z')p_k(z',t) \right) p_p(z,t)dz \qquad (4)$$

here, $b_p$ is the growth rate independent of competitive interactions and predation pressure; $\alpha_{\mathrm{pj}}^P$ captures the competitive effect of basal species $j$ on basal species $p$ ; $b(z,z')$ is the Gaussian predation function that captures the likelihood of an individual with trait $z$ of basal species being predated by an individual with trait $z'$ from the consumers with trait distribution given by $p_k(z',t)$. The more similar the traits are the more likely is a consumer able to successfully predate on an individual from basal species with trait $z$. Thus population dynamics can be written by substituting the above equation in equation 1 of the main-text :

$$\frac{dN_p}{dt} = N_p(t) \left( b_p - \sum_{j\epsilon \ basal \ sp.} \alpha_{\mathrm{pj}}^P(t)\, N_j(t) - \sum_{k\epsilon \ consumers} C_k(t)\beta_{pk}(u_p,u_k) \right) \qquad (5)$$

where, $C_k$ is the density of a consumer species $k$, $\beta_{pk}$ captures the trait-mediated predation of consumer species $k$ on basal species $p$ ;$\alpha_{\mathrm{pj}}^P$ captures both intra and interspecific competiton among the basal species that is independent of mean trait values. $\beta_{pk}(u_p,u_k) = \int\int b(z,z')p_k(z',t)dz'p_p(z,t)dz = \frac{b}{\sqrt{2\sigma_k^2+2\sigma_p^2+b^2}} \exp^{-\frac{(u_k-u_p)^2}{2\sigma_k^2+2\sigma_p^2+b^2}}$; where $b$ captures the strength of predator-prey interaction between the basal species and the consumers.

Similarly, for the consumer species population dynamics can be written as:

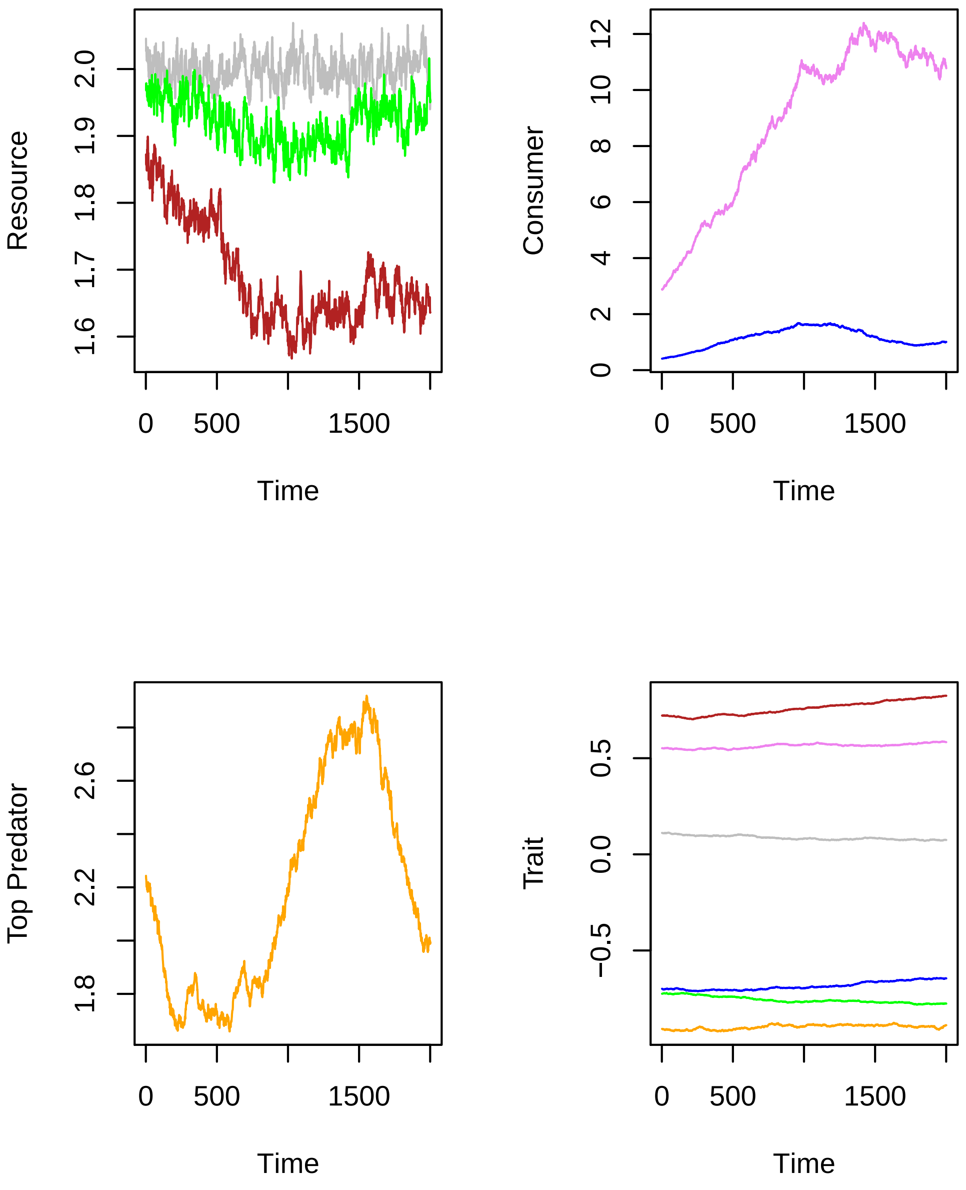


Figure S3: Time series of abundance and trait of all six species with the parameter values mentioned in the appendix for the predator-prey community.

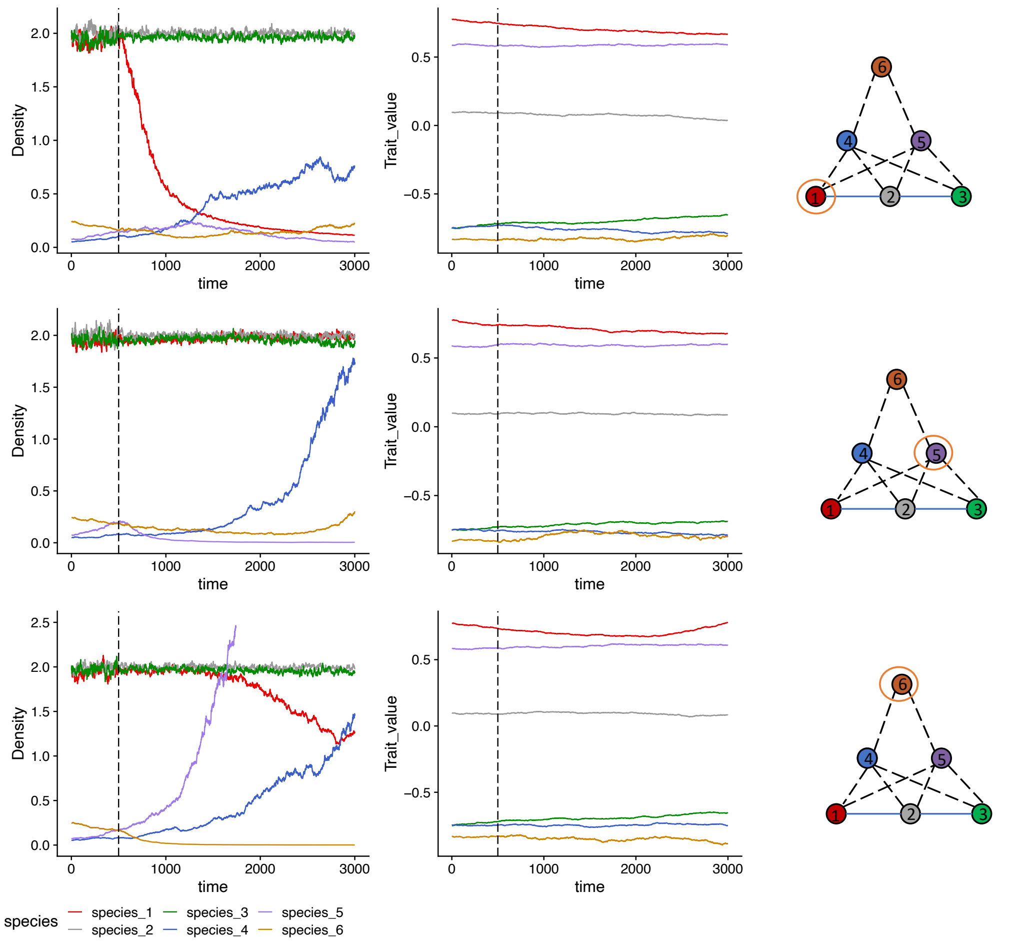


Figure S4: Time series of abundance and trait of all six species with the parameter values mentioned in the appendix for the predator-prey community. Dashed vertical line indicates the timepoint of species forcing. In the community motif circled species index number is the one being forced to collapse while rest of the species in the community remains unperturbed.

$$\frac{dC_p}{dt} = C_p\left(d_C - \sum_{j\epsilon\ consumers} C_j(t)a^C_{pj}(t) + \sum_{k\ \epsilon\ basal\ sp.} \beta_{pk}(u_p, u_k)(t)N_k(t) - E_{pl}(t)P_l(t)\right) \tag{6}$$

where, $d_C$ is the mortality rate of the consumer species; $\beta_{pk}(u_p, u_k)$ captures fitness gain from trait-mediated predation of basal species $k$; $E_{pl}$ captures the predation from the top predator on the consumers (species 4, 5)in the predator-prey foodweb community. $E_{pl} = \frac{b}{\sqrt{2\sigma_p^2+2\sigma_l^2+b^2}} \exp^{-\frac{(u_p-u_l)^2}{2\sigma_p^2+2\sigma_l^2+b^2}}$ is also a Gaussian function similar to the one above.

And finally the dynamics of the top predator can be written as:

$$\frac{dP_p}{dt} = P_p\left(d_p - a_{pp}P_p + \sum_{l\ \epsilon\ consumers} E_{pl}(t)C_l(t)\right) \tag{7}$$

For all the basal species, $b_p$ was kept at 1; for the consumers and the top predators,$d_C$ and $d_p$ were mortality rates drawn in accordance with Petchey et. al 2008. Predator-prey interaction strength of consumers on the basal was kept at $b = 0.008$ and for top-predators on consumers was also kept at $b = 0.008$.

We ran simulations for 5000 time points with varying mean traits values for each species in the community from random uniform distribution $[-1, 1]$ until we found a set of mean trait values that led to feasible species densities. We used these mean trait values as the starting point of our simulations for population collapses in the mixed interaction community. Initial abundances of the all the species were kept at 1. Strength of predator-prey interaction both between the basal species and the consumers, and between the consumers and the top predator were kept at 0.01. Species trait variances for the competition community were randomly assigned for each replicate simulation of population collapse from a random uniform distribution $[0.005, 0.02]$. At each time step of population dynamics we added Gaussian white noise with mean zero and variance of 0.5. Mean trait dynamics $u_i$ of a species $i$ followed from equation 6.2 of the main-text.

### 3. Effective potential curves

#### 3.1. Mutualistic interaction

For mutualistic interactions, the effective potential can be written as:

$$V_m(N_i) = -\int \left( b_i^m(u_i) - \sum_{j=1}^{N} \alpha_{ij}^m N_j + \sum_{K=1}^{A} A_k \gamma_{ik} \right) N_i dN_i$$

which gives :

$$V_m(N_i) = -b_i^m(u_i)\frac{N_i^2}{2} - \alpha_{ii}^m \frac{N_i^3}{3} - \sum_{i \neq j}^{N} \alpha_{ij}^m N_j \frac{N_i^2}{2} + \sum_{K=1}^{A} A_k \gamma_{ik} \frac{N_i^3}{3} \tag{8}$$

Specifically, we estimate $V_m(N_i)$ just before environmental forcing starts i.e., at $t = 800$ time point. At that time point, we estimated three quantities that accurately characterize the potential curves : a) slope of the potential curve which quantifies how stable the one-dimensional system is; b) depth of the potential curve given as: $min(V_m(N_i) - max(V_m(N_i))$ , which quantifies how resilient the system is; and c) scaled potential width, which is the distance between the tipping point and the lowest point of the potential. Potential width quantifies the amount of perturbation a one-dimensional system such as a population can withstand before transitioning to another state [1]. We quantify these three metrics just before the start of environmental forcing for each replicate population collapses (n=100) and for each of the community modules, and try to evaluate whether different interaction types modify these stability-resilience metrics and in turn influence the utility of EWS of population collapses.

#### 3.2. Predator-prey interactions

For the basal species, the effective potential can be written as at time point $t$ as:

$$V_b(N_p) = -\int \left( b_p(u_p) - \sum_{j \ \epsilon \ basal \ sp} \alpha_{pj}^p N_j - \sum_{k \ \epsilon \ consumers} C_k \beta_{pk} \right) N_p dN_p$$

which gives:

$$V_b(N_p) = -\left( b_p(u_p)\frac{N_p^2}{2} - \alpha_{pp}^p N_p^3/3 - \sum_{p \neq j} \alpha_{pj} N_j \frac{N_p^2}{2} - \sum_{k \ \epsilon \ consumers} C_k \beta_{pk} \frac{N_p^2}{2} \right) \tag{9}$$

And for the consumer species 4 and 5,

$$V_c(C_i) = -(d_c \frac{C_i^2}{2} - a_{ii}^c C_i^3/3 - \sum_{i \neq k} a_{ik} C_k \frac{C_i^2}{2} + \sum_{k \ \epsilon \ basalsp.} N_k \beta_{ik} \frac{C_i^2}{2} - E_{il} P_l \frac{C_i^2}{2}) \tag{10}$$

And finally for the top predator (species 6):

$$V_p = -(d_P \frac{P^2}{2} - a_{pp}^P \frac{P}{3} 3 + \sum_{k \ \epsilon \ consumers} E_k C_k \frac{P^2}{2} \tag{11}$$

Specifically, we estimate the effective potentials of the basal species,$V_b(N_p)$ consumers $V_c(C_i)$ and the top predators $V_p$ just before environmental forcing started. At that time point, we estimated three quantities that accurately characterize the potential curves : a) slope of the potential curve which quantifies how stable the one-dimensional system is; b) depth of the potential curve, which quantifies how resilient the system is; and c) scaled potential width, which is the distance between the tipping point and the lowest point of the potential. Potential width quantifies the amount of perturbation a one-dimensional system such as a population can withstand before transitioning to another state [1]. We quantify these three metrics (see Fig. 1) just before the start of environmental forcing for each replicate population collapses (n=100) and for each of the community modules, and try to evaluate whether different interaction types modify these stability-resilience metrics and in turn influence the utility of EWS of population collapses.

### References

[1] Nolting, B.C. & Abbott, K.C. (2015) Balls, cups, and quasi-potentials: quantifying stability in stochastic systems. *Ecology* **97**, 15–1047.1.
